# Vascular Smooth Muscle Cell Mechanotransduction Through Serum and Glucocorticoid Inducible Kinase -1 Promotes Interleukin-6 Production and Macrophage Accumulation in Murine Hypertension

**DOI:** 10.1101/2023.03.03.530966

**Authors:** Mario Figueroa, SarahRose Hall, Victoria Mattia, Alex Mendoza, Adam Brown, Ying Xiong, Rupak Mukherjee, Jeffrey A. Jones, William Richardson, Jean Marie Ruddy

**Author notes:** Corresponding Author Jean Marie Ruddy, MD FACS Associate Professor of Surgery Division of Vascular Surgery Medical University of South Carolina 30 Courtenay Drive, MSC 295 Ste 654, Charleston, SC 29425.

## Abstract

**Objective:** The objective of this investigation was to demonstrate that *in vivo* induction of hypertension (HTN) and *in vitro* cyclic stretch of aortic VSMCs can cause SGK-1-dependent production of cytokines to promote macrophage accumulation as agents of vascular remodeling.

**Methods:** HTN was induced in C57Bl/6 mice with AngiotensinII (AngII) infusion (1.46mg/kg/day x 21 days) with or without systemic infusion of EMD638683 (2.5mg/kg/day x 21 days), a selective SGK-1 inhibitor. Systolic blood pressure (SBP) was recorded on days 0 and 21. At terminal study, abdominal aortas were harvested to quantify SGK-1 activity (pSGK-1:SGK-1) by immunoblot. Additional replicates were digested and analyzed by flow cytometry for abundance of CD11b^+^/F4-80^+^ cells (macrophages). Plasma was analyzed by ELISA to quantify IL-6 and MCP-1. Aortic VSMCs from wild-type (WT) mice were subjected to 12% biaxial cyclic stretch for 3 or 12 hours +/- EMD638683 (10μM) and +/- SGK-1siRNA with subsequent QPCR for IL-6 and MCP-1 expression. Culture media was analyzed by ELISA for IL-6 and MCP-1. Aortic VSMCs from SGK-1^flox+/+^ mice were transfected with Cre-Adenovirus to knockout SGK-1 (SGK-1KO VSMCs) and underwent parallel tension experimentation. Computational modeling was employed to simulate VSMC signaling due to mechanical strain and AngII. Statistical analysis included ANOVA with significance at p<0.05.

**Results:** SGK-1 activity (pSGK-1:SGK-1) was upregulated in the abdominal aorta of mice with HTN and significantly reduced by treatment with EMD638683. Concurrently, increased CD11b^+^/F4-80^+^ cells and plasma IL-6 levels in the HTN group and reduction with EMD638683 was observed. This mirrored the increased abundance of IL-6 in media from Stretch WT VSMCs, and attenuation of the effect with EMD638683. Treating WT VSMCs with SGK-1siRNA likewise inhibited IL-6 expression. MCP-1 also demonstrated increased expression and secretion into the media in WT VSMCs with Stretch. Further supporting the integral role of mechanical signaling through SGK-1, target gene expression and cytokine secretion was unchanged in SGK-1KO VSMCs with Stretch, and computer modeling confirmed SGK-1 as an intersecting node of signaling due to mechanical strain and AngII. In summation, this data suggests a biomechanical link between aortic VSMC mechanotransduction and cytokine production to promote macrophage accumulation, mediated in-part by SGK-1 activation.

**Conclusion:** Mechanotransduction through SGK-1 is instrumental in pro-inflammatory cytokine production and aortic macrophage accumulation in systemic HTN, therefore further investigation into targeting this kinase may present opportunities to modulate hypertensive vascular remodeling.

## Introduction

In hypertension (HTN), the aorta is subject to shear stress and elevated wall tension that has been associated with dysfunctional remodeling, stiffening, and increased cardiovascular mortality.^1,2^ Likewise, human samples as well as animal models have demonstrated leukocyte infiltration in hypertensive vascular disease.^3^ The question of whether the wall stress caused by these hemodynamic forces is a product of the inflammatory infiltrate or if it contributes to the propagation of detrimental inflammatory signaling cascades remains unanswered. This laboratory has previously demonstrated that Interleukin-6 (IL-6) and monocyte chemoattractant protein-1 (MCP-1) expression was elevated in murine aortic VSMCs under conditions of biaxial stretch and interestingly, transcriptional activity was not enhanced by concurrent treatment with the vasoactive peptide AngiogensinII (AngII).^4^ Targeted interruption of this mechanical signal transduction may represent an opportunity to quench dysfunctional vascular remodeling.

With a focus on identifying the tension-induced pathway implicated in propagating hypertensive pro-inflammatory signaling, it was noted that the serum and glucocorticoid-inducible kinase-1 has already demonstrated mechanosensitivity in other vascular beds such as vein bypass grafts.^5^ Furthermore, in pulmonary HTN, blockade of SGK-1 improved pressure differentials and reduced macrophage infiltration.^6,7^ Current evidence relating SGK-1 activity to systemic HTN has focused on renal ion channel production and function in salt-induced models,^8^ such that knockout animals were protected from blood pressure elevation and consequential inflammation.^9,10^ Regarding the intrinsic aortic cell response to tension, utilization of the AngII-induced murine HTN model has advantageous clinical translation, and infusion of that biologic stressor has likewise demonstrated the aorta as a specific target of damage and inflammation under conditions of HTN.^11^ SGK-1 activity within the aortic wall of AngII-induced HTN has not been described and represents an opportunity to investigate the integrated biologic and mechanical signaling pathways in HTN. Accordingly, this study focused on further specifying the role of SGK-1 in hypertensive aortic inflammation by hypothesizing that SGK-1 activity is upregulated under conditions of elevated tension to promote pro-inflammatory cytokine production and macrophage accumulation which may lead to dysfunctional aortic remodeling.

## Methods

### Animal Care and Use

All animal care and surgical procedures were approved by the Medical University of South Carolina Institutional Animal Care and Use Committee (AR#2020-01502). C57Bl/6 mice were purchased from Jackson Laboratories (Bar Harbor, ME) and local breeding colony was established to support experimentation. SGK-1^flox+/flox+^ mice on a C57Bl/6 background were obtained from the Fejes-Toth laboratory (Dartmouth University).^12^ A local breeding colony was established.

### AngII-induced HTN and SGK-1 Inhibition with EMD638683

AngII was dissolved in saline and loaded at concentrations to deliver 1.46 mg/kg/day for 21 days via mini pump. Prior experimentation in this laboratory utilizing this model of murine HTN has confirmed that AngII infusion in normo-lipidemic, wild-type mice of this age and fed regular chow did not develop aneurysms.^13^ EMD638683 (MedChemExpress, Monmouth Junction, NJ), a selective inhibitor of SGK-1 with no identified alternate targets, was dissolved in DMSO and stored in aliquots at 4°C. Aliquots were further diluted to achieve 2.5 mg/kg/day delivery for 21 days via mini pump.

Following induction of anesthesia with 3% isoflurane and subcutaneous injection of 0.05 mg/kg buprenorphine, C57Bl/6 mice (12-20 weeks of age) underwent subcutaneous left flank implantation of loaded Alzet osmotic mini pump (model 1004; Durect Corporation, Cupertino, CA). C57Bl/6 mice received either an AngII infusion (C57Bl/6+AngII), EMD638683 infusion (C57Bl/6+EMD), or both treatments (C57Bl/6+AngII+EMD) simultaneously. Following implantation, the incision was closed and sutures were removed 14 days after surgery.

### Blood Pressure Measurement

Mouse blood pressures were measured on Day 0 and Day 21 of treatment. CODA8 tail-cuff method (Kent Scientific, Torrington, CT) was utilized as previously described.^13,14^ Briefly, mice were allowed 10 minutes of acclimation to the chamber. A dark and warm environment was maintained throughout data collection with a heating pad and towel cloak.

### Tissue Harvest and Processing

All terminal procedures were conducted on Day 21. Following induction of 3% isoflurane, laparotomy was performed, small bowel was mobilized, and the aorta was exposed from the renal vein to the bifurcation. Terminal aortic diameter measurements were taken via calibrated digital microscopy. Infrarenal aortas were harvested and flash frozen in a liquid nitrogen slurry. Blood from the left ventricle was collected with an EDTA treated syringe, and centrifuged at 2500xg for 15 minutes at 4°C. Plasma was harvested and samples were stored in -80°C until further analysis.

### Immunoblot Analysis

Relative abundance of SGK-1, pSGK-1, the mature murine macrophage marker F4/80, and α-tubulin were determined by immunoblotting. Harvested aortic tissue was homogenized in cell lysis buffer and protease inhibitor cocktail via bead homogenization. Following homogenization, 20 μg of protein from each abdominal aortic tissue sample, as determined by the Pierce BCA total protein assay (23225; Thermo Fisher Scientific, Waltham, MA), was fractionated on a 10% polyacrylamide gel by electrophoresis. The proteins were transferred to nitrocellulose membranes (0.45 μm; Bio-Rad, Hercules, CA) and incubated in antisera specific for SGK-1 (ab32374, AbCam, Waltham, MA; 1:1000), pSGK-1 (ab55281, AbCam, Waltham, MA; 1:1000), F4/80 (ab6640, AbCam, Waltham, MA; 1:1000), and α-tubulin (ab7291, AbCam, Waltham, MA 1:1000) in 0.1% Tween/TBS. A secondary peroxidase-conjugated antibody was applied (1:5000; 0.1% TBST). Signals were detected with a chemiluminescent substrate (Western Lighting Chemiluminescence Reagent Plus; PerkinElmer, San Jose, CA) and recorded on film. Between each protein analysis, membranes were stripped using Restore Stripping Buffer (Thermo Fisher Scientific, Waltham, MA). Band intensity was quantified using ImageJ 53C software (National Institute of Health, USA) and data was reported as a fold change from normotensive control.

### Flow Cytometry

In mice designated for flow cytometric analysis, freshly harvested infrarenal aortas were collected in Aortic Enzyme Digest Solution (125 U/mL Collagenase type XI, 450 U/mL Collagenase type I, 60 U/mL DNase-1, and 60 U/mL Hyaluronidase type-1-s in 2.5 mL of Dulbecco’s PBS) and incubated at 37°C in 5% CO_2_ for 2 hours. After digestion, the cells were passed through a 35μm cell strainer snap cap (352235; Corning Life Sciences, Durham, NC), rinsed with 1 mL BD Pharmingen Stain Buffer (554656; BD Biosciences. San Diego, CA), and centrifuged at 300xg at 4°C. The cell pellet was resuspended in 1 mL FACS Buffer, transferred to a new 4 mL snap cap Falcon tube (149592A; Corning Life Sciences, Durham, NC), and centrifuged at 300xg at 4°C. Cells were blocked with mouse Fc receptor block (Miltenyi Biotec, San Diego, CA) and stained with conjugated primary antibodies: CD11b-FITC (clone M1/70, BD Biosciences, San Jose, CA) and F4/80-BV421 (clone BM8, Biolegends, San Diego, CA) for 20 min at 4°C. Cells were then washed, centrifuged at 300xg for 10 min, and resuspended in BD Pharmingen Stain buffer. Cells were fixed by adding 4% Formalin/PBS and incubated for 15 min at 23°C. Cells were then washed in BD Pharmingen Stain buffer and centrifuged at 300xg, and resuspended in BD Pharmingen Stain buffer.

Samples were processed on a CytoFLEX Flow Cytometer (Beckman Coulter) equipped with CytExpert Software (Beckman Coulter). An average of 100,000 events were recorded for each sample. Analysis of the results was performed using CytExpert software (Version 2.4.0.28, Beckman Coulter). The gating included cell size selection, followed by single cell isolation through side-scatter (Supplement Figure 1). The remaining events were gated for concurrent staining with myeloid marker CD11b^+^ and the mature macrophage marker F4/80^+^.

### Blocking SGK-1 in VSMC Culture

Primary VSMC lines from C57Bl/6 and SGK-1^flox+/+^ aortic biopsies were established using an accepted outgrowth technique as previously described.^15^ Isolated VSMCs were maintained in SMC specific growth media with added supplement pack (SMC Growth Medium 2; C-22062, PromoCell, Heidelberg, Germany) at 37°C in 5% CO_2_. For SGK-1^flox+/+^ VSMCs that had reached confluency, a kanamycin-resistant Cre-adenovirus (SignaGen, Fredrick, MD) was utilized to mute transcription of SGK-1 in that subset population. Virus containing media at 100 MOI was diluted as recommended and administered to SGK-1^flox+/+^ VSMCs. Following 24 hours of infection, cells were treated with kanamycin (Millipore Sigma, Burlington, MA) to remove any non-transfected VSMCs. These were subsequently referred to as SGK-1KO VSMCs.

Confluent C57Bl/6, SGK-1^flox+/+^, and SGK-1KO VSMCs from culture passages 2-10 were seeded at a density of 5000 cells per cm^2^ into amino coated Bioflex-6 well plates (BF-3001A; Flexcell International Corporation, Burlington NC) and allowed to adhere overnight. Complete SMC medium was replaced with serum free SMC basal media (SMC Basal Medium 2; C-22262, PromoCell. Heidelberg, Germany) and allowed to rest for 6 hours at 37°C in 5% CO_2._ VSMCs were treated with +/- EMD638683 (SGK-1-specific inhibitor; 10μM) and allowed to rest at 37°C in 5% CO_2_ for 1 hour. Following this incubation, plates were held under Static conditions or subjected to 12% biaxial cyclic Stretch for 3 hours or 12 hours utilizing the Flexcell culture system (Flexcell International Corporation, Burlington, NC). Cells were harvested to quantify expression of IL-6 and MCP-1 by QPCR.

Additionally, siRNA transfection technique was utilized to silence SGK-1 activity in wild-type aortic VSMC. C57Bl/6 cells were seeded overnight in 6-well plates. An SGK-1 specific siRNA and Lipofectamine RNAiMAX cocktail were diluted in culture medium as recommended, as was a silencer select RNA negative control and Lipofectamine RNAiMAX cocktail (LifeTechnologies, Carlesbad, Ca). Cells were allowed 48 hours incubation in the either the SGK-1 specific siRNA transfection media, silencer select transfection media, or control media. Following transfection, cells were lifted and seeded (5000 cells/cm^2^) overnight on amino coated Bioflex-6well plate (Flexcell International Corporation, Burlington NC) and subjected to Static or Stretch conditions +/- EMD638683 as described above. Cells were harvested to quantify expression of IL-6 and MCP-1.

### Quantitative Polymerase Chain Reaction (QPCR)

Following treatment on the Flexcell system, total RNA was extracted using TRIzol Reagent (15596026, Thermo Fisher Scientific, Waltham, MA). One microgram of total RNA, determined by NanoDrop 2000 (Thermo Fisher Scientific, Waltham, MA), was reverse transcribed and converted to complementary DNA (cDNA) using the iScript cDNA synthesis kit (1708891, Bio-Rad, Hercules, CA). Each cDNA sample was amplified with messenger RNA (mRNA) specific TaqMan Gene Expression Assays (IL-6, Mm00446190_m1; MCP-1, Mm00441242_m1; GAPDH, Mm99999915_g1; Thermo Fisher Scientific, Waltham, MA) on a CFX-96 real-time polymerase chain reaction (PCR) machine (Bio-Rad). The relative expression of each mRNA was calculated and normalized to the expression of glyceraldehyde 3-phosphate dehydrogenase (GAPDH). Expression values were calculated as 2^−ΔCT^, in which the change in cycle threshold was defined as normalized gene expression. Values were then reported as a fold change compared to expression in the respective Static VSMCs.

### ELISA

Enzyme- linked immunosorbent assay (ELISA) was used to quantify the concentration of IL-6 (ab100712; Abcam, Waltham, MA) and MCP-1 (ab208979; Abcam, Waltham, MA) in Flexcell treatment conditioned media and murine plasma. Collected cell media was first concentrated using a 10K Centrifugal Filter (UFC201024; Millipore Sigma, Carrigtwohill, CO) and centrifuged at 4000xg for 30 min at 4°C. The sample recovery was plated in the respective 96-well strip plates. Standard dilution and reagents were prepared in accordance to manufacturer’s instructions. Absorbance at 450 nm was measured using a SpectraMax M3 microplate absorbance reading (Molecular Devices) with SoftMax Pro (Version 7.1) software. Abundance values were then reported as a fold change from the wild-type condition.

### Computational Modeling

In order to better elucidate the role of SGK-1 within the broader cell signaling pathways connecting mechanical stimulation and IL-6 production, we adapted a previously published computational model of VSMC signaling networks.^16–18^ Signaling proteins and small molecules included in network model represent the most well-established pathways connecting AngII and mechanical strain inputs to changes in cell expression related to IL-6 expression, as well as matrix synthesis (e.g., collagens), matrix degradation (e.g., MMPs), and cell contractility (e.g., acto-myosin proteins). These pathways were manually reconstructed based on extensive mechanistic studies in the literature, and the resulting network spans 79 molecules interconnected by 123 reactions including the well-established signaling drivers Akt, Rho, Smads, JNK, ERK, MAPK, p38, mTOR, PI3K, and others. We adapted the previous network by adding SGK-1 reactions supported by previous literature.^19^ The full list of integrated species and reactions are available as Supplement Tables 1 and 2.

To simulate the effects of input stimuli on downstream nodes, each connection in the model is simplified as a normalized Hill differential equation according to logic-based gates (AND, OR relationships, Equations 1-2):

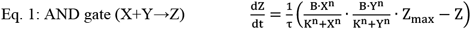

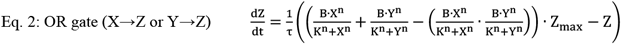

where X, Y, Z are node activation levels, τ is a reaction time constant, and n is the classic Hill coefficient that captures the nonlinearity of each reaction’s dose-response curve. B, K are additional parameters describing dose-response features and determined by n and the half-maximal activation concentration, EC_50_, according to the Equations 3-4:

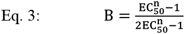

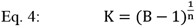

The system of equations is solved using MATLAB code driven by the open-source, Netflux user interface [https://github.com/saucermanlab/Netflux]. ^20^

Given specific input inhibition or stimulation, the model calculates resulting changes in the expression of primary outputs (IL-6, ECM, MMPs, etc.). This same logic-based Hill equation approach has been applied across a variety of cell types including VSMCs, vascular adventitial fibroblasts, cardiac fibroblasts, valve interstitial cells, macrophages, and cardiomyocytes.^16–18,20–27^ Herein, we represented the *in vitro* experimental conditions by running a baseline control simulation, a heightened mechanical strain simulation, and a strain + SGK-1-inhibition simulation. We represented the *in vivo* experimental conditions by running a baseline control simulation, a heightened mechanical strain + AngII simulation, and a strain + AngII + SGK-1-inhibition simulation.

### Statistical Analysis

The percent increase in abdominal AoD was calculated as ([terminal AoD – baseline AoD/baseline AoD] x 100 +100) for each individual mouse. Therefore, the baseline value for each mouse is 100% and reported results represent the increase in diameter above baseline. The pSGK-1/SGK-1 ratio was calculated for each aortic sample and the fold change established relative to the C57Bl/6 control mice which represent normal aortic physiology. Flow cytometric analysis quantified the CD11b^+^/F480^+^ cells as a percentage of the total cells within the harvested abdominal aortic tissue. QPCR and ELISA values were represented as fold change from baseline. In each data set, the groups were compared by ANOVA and *post hoc* mean separation was performed using the Sidak correction for multiple comparisons. Statistical tests were performed using Stata (v17.0, College Station, TX) with significance established at p<0.05.

## Results

In C57Bl/6 wild-type mice, AngII infusion caused a significant increase in systolic blood pressure (SBP) at 21 days which was not impacted by EMD treatment (Figure 1). Consistent with prior utilization of this model in wild-type mice, at terminal procedure neither aortic dilation nor dissection was apparent (data not shown). Abdominal aortic tissue from C57Bl/6+AngII mice demonstrated significant elevations in SGK-1 activity and macrophage infiltration (p<0.05 vs C57Bl/6) which were effectively inhibited by EMD infusion (p<0.05 vs C57Bl/6+AngII; Figures 2A and 2B). Although plasma levels of MCP-1 were not detectable, IL-6 was increased >1.5 fold (p<0.05 vs C57Bl/6) and significantly reduced by EMD (p<0.05 vs C57Bl/6+AngII; Figure 2C). It has been well documented that even patients with controlled HTN can develop end organ damage, therefore these results suggest that blocking SGK-1 activity and downstream inflammatory markers may represent a complementary treatment paradigm to combat hypertensive aortic remodeling.

**Figure 1.**
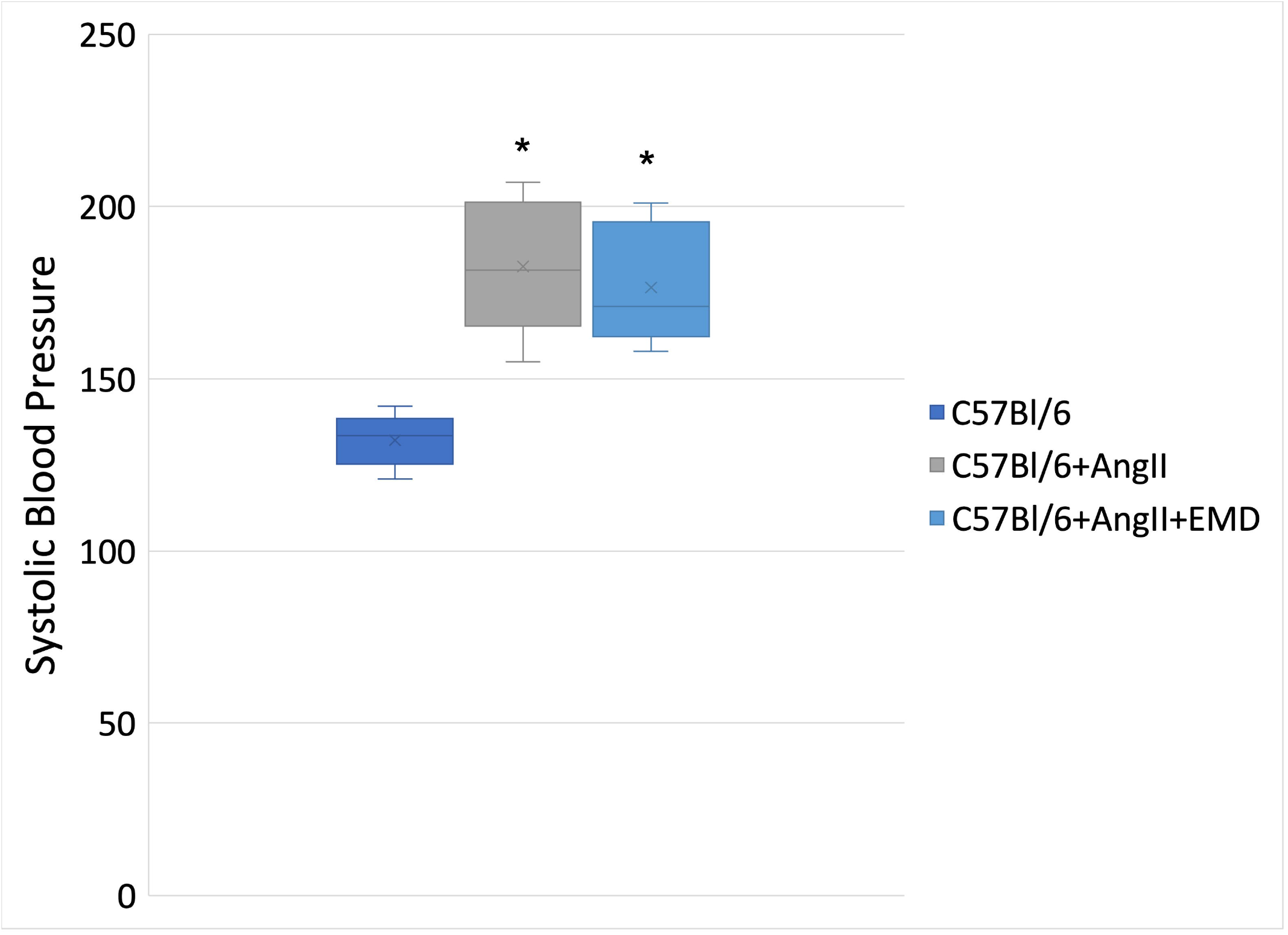
Systolic blood pressure as measured by tail cuff. *p<0.05 vs Day 0

**Figure 2.**
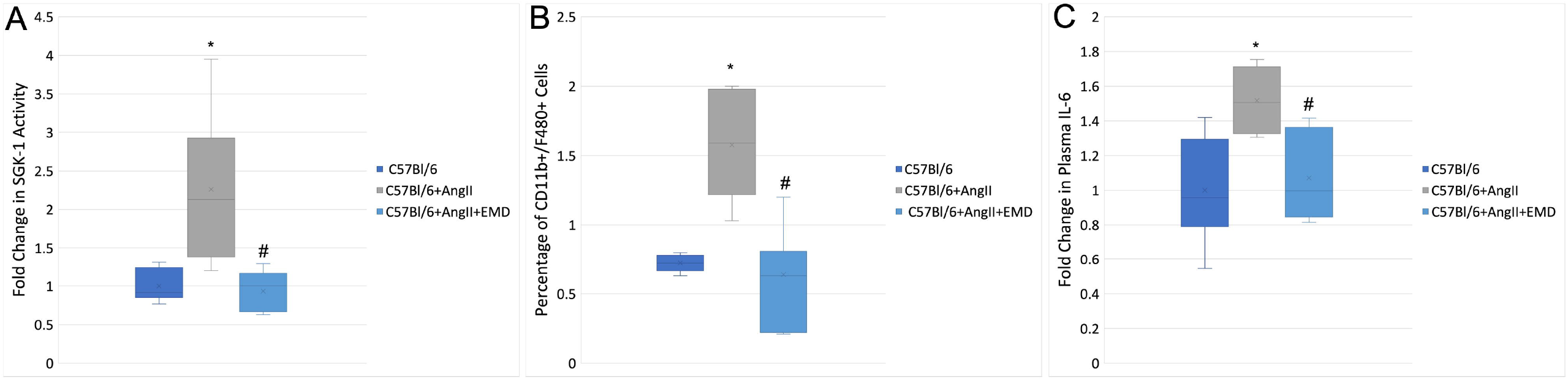
(A) Fold change in SGK-1 activity (pSGK-1/SGK-1) in aortic homogenate at Day 21. (B) Flow cytometric quantification of macrophages in abdominal aorta on Day 21. (C) Quantification of IL-6 in plasma of experimental mice at Day 21 represented as a fold change from the control. *p<0.05 vs C57Bl/6; #p<0.05 vs C57Bl/6+AngII

When aortic VSMCs were harvested from C57Bl/6 mice, 3 hours of cyclic biaxial Stretch led to significant increase in IL-6 and MCP-1 expression (p<0.05 vs Static, Figures 3A and 4A), which was inhibited by treatment with EMD or SGK-1siRNA (p<0.05 vs Stretch) but dual therapy did not have an additive effect. Aortic VSMCs harvested from SGK-1^flox+/+^ mice were treated with Cre-Adenovirus to generate SGK-1KO VSMCs which underwent 12 hours of Stretch compared to C57Bl/6 wild-type VSMCs. IL-6 expression in WT VSMCs and secretion of IL-6 into the culture media followed the same pattern of Stretch-induced elevation with EMD inhibition, but no response was observed for SGK-1KO VSMCs (Figure 3B and 3C). Interestingly, with 12 hours of Stretch, MCP-1 expression in C57Bl/6 WT cells trended back to baseline Static levels to suggest a negative feedback system to quench transcription. Translational activity continued, however, and the MCP-1 protein abundance in the culture media was significantly elevated (p<0.05 vs WT Static; Figures 4B and 4C). In the SGK-1KO VSMCs, the effect of that negative feedback actually led to decreased MCP-1 expression with prolonged Stretch or EMD treatment. Integrating this isolated VSMC data with the murine AngII-induced HTN model may support that mechanical activation of SGK-1 in aortic VSMCs can promote IL-6 production to raise serum levels as well as local tissue MCP-1 levels to contribute to macrophage infiltration that propagates dysfunctional remodeling.

**Figure 3.**
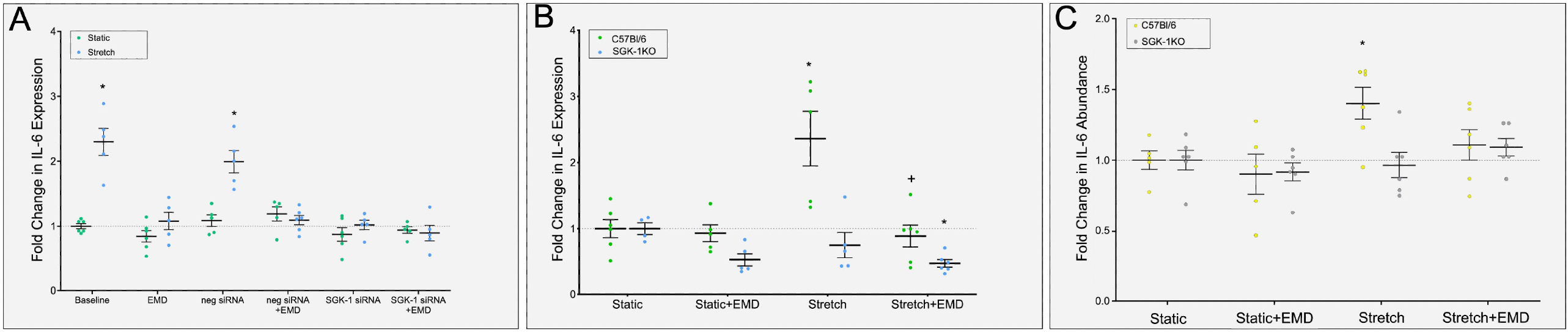
(A) IL-6 expression in C57Bl/6 WT VSMCs treated +/- Stretch, +/- siRNA, and +/- EMD for 3 hours. (B) IL-6 expression in C57Bl/6 WT and SGK-1KO VSMCs treated for 12 hours +/- Stretch and +/- EMD. (C) IL-6 abundance in culture media from C57Bl/6 WT and SGK-1KO VSMCs treated for 12 hours +/- Stretch and +/- EMD. *p<0.05 vs WT Static, +p<0.05 vs WT Stretch

**Figure 4.**
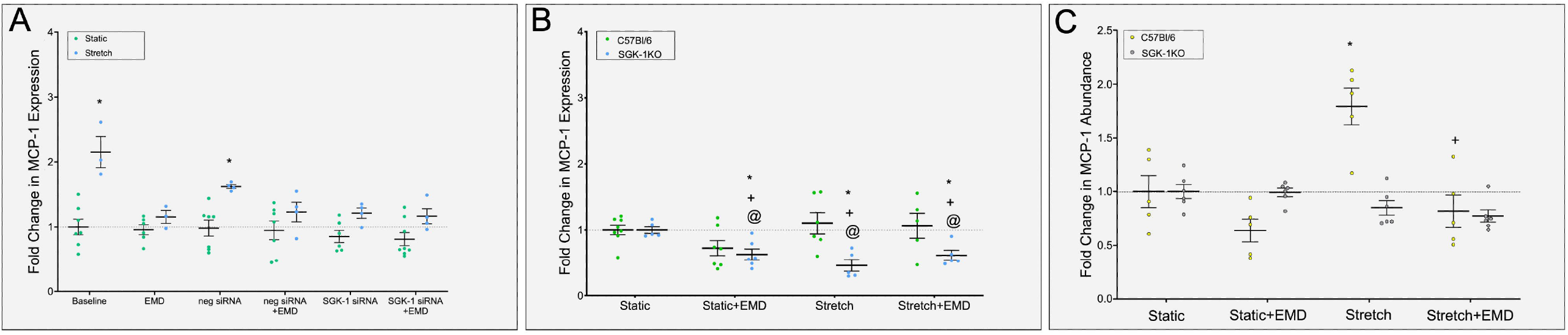
(A) MCP-1 expression in C57Bl/6 WT VSMCs treated +/- Stretch, +/- siRNA, and +/- EMD for 3 hours. (B) MCP-1 expression in C57Bl/6 WT and SGK-1KO VSMCs treated for 12 hours +/- Stretch and +/- EMD. (C) MCP-1 abundance in culture media from C57Bl/6 WT and SGK-1KO VSMCs treated for 12 hours +/- Stretch and +/- EMD. *p<0.05 vs WT Static, +p<0.05 vs WT Stretch, @p<0.05 vs SGK-1KO Static

Prior experimentation had indicated that adding AngII treatment to mechanical stretch in murine aortic VSMCs did not amplify IL-6 expression beyond that of stretch alone,^28^ therefore dual stimulation concurrent to SGK-1 inhibition was unlikely to strengthen the clinical relevance of this project. Instead, we chose to employ computational modeling of VSMC signaling to further evaluate SGK-1 as a major contributor to tension-induced pro-inflammatory signaling in HTN. Signaling proteins and small molecules included in the VSMC network model (Figure 5A) represent the most well-established pathways connecting AngII and mechanical strain inputs to changes in cell expression related to inflammation (e.g., IL-6), matrix synthesis (e.g., collagens), degradation (e.g., MMPs), and cell contractility (e.g., acto-myosin proteins). As noted above in the Methods, these pathways were manually reconstructed based on extensive mechanistic studies in the literature. Importantly, the network has included major feedback loops driven by IL-6, and SGK-1 represented a central hub connecting both AngII and mechanical strain inputs to inflammatory, synthetic, degradation, and contractility outputs. Several pertinent positive (Figure 5B) and negative (Figure 5C) feedback loops were observed. Notably, inhibiting SGK1 could interrupt at least three positive feedback loops that may drive IL-6 expression in aortic VSMCs:

**Figure 5.**
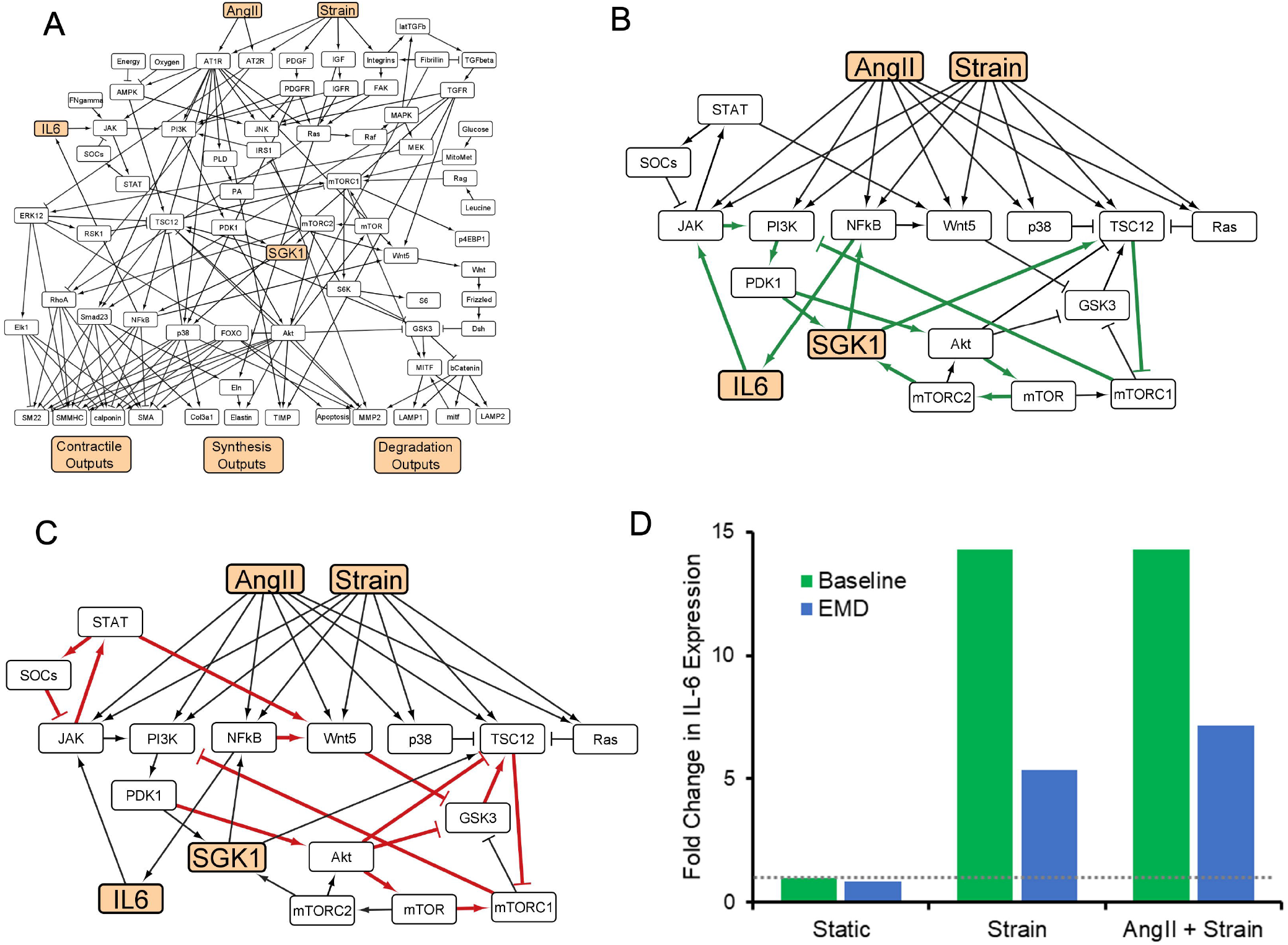
(A) A network model of SMC signaling enabling quantitative, differential equation-based predictions of cell expression responses to AngII and mechanical strain. (B-C) Further, a simplified model topology identified negative feedback loops (B) and positive feedback loops (C) that contribute to the complexity of IL-6 regulation. (D) Model prediction that inhibiting SGK-1 activity with EMD638683 results in a substantial reduction of IL-6 expression in VSMCs under mechanical and AngII treatments.

IL-6→JAK→PI3K→PDK1→***SGK-1***→NFkB→IL-6

IL-6→JAK→PI3K→PDK1→Akt→mTOR→mTORC2→***SGK-1***→NFkB→IL-6

***SGK-1***→TSC12→mTORC2→PI3K→PDK1→***SGK-1***

Concordantly, the computational simulations were able to replicate our *in vitro* findings that inhibition of SGK-1 activity led to a decrease in expression of IL-6 (Figure 5D). The integration of this *in vivo, in vitro*, and computational data strongly supports SGK-1 as a major mechanosensitive kinase in hypertensive vascular inflammation that may be targeted to reduce pathologic remodeling.

## Discussion

Moving beyond the role of SGK-1 in renal salt secretion, this project has demonstrated that in HTN, the mechanical activation of SGK-1 in aortic VSMCs can promote inflammation known to facilitate vascular remodeling. In the AngII-induced HTN model, inhibiting SGK-1 with continuous infusion of EMD638683 did not effect SBP but significantly reduced plasma IL-6 as well as macrophage infiltration to demonstrate that despite the ongoing mechanical stimulus, intracellular SGK-1 inhibition reduced inflammatory signaling. The influence of mechanical activation of SGK-1 on cytokine production was further established in harvested murine aortic VSMCs wherein cyclic stretch fostered IL-6 and MCP-1 expression as well as protein secretion, but this was inhibited with pharmacologic or genetic blockade of SGK-1 activity. Furthermore, computational modeling of VSMC signaling due to mechanical strain and/or AngII stimulation identified pertinent positive feedback loops centered on SGK-1 to upregulate IL-6 expression. Considering the micro- and macrovascular impact of pathologic hypertensive remodeling, targeting SGK-1 to reduce pro-inflammatory signaling may represent a vital adjunct to standard anti-hypertensive therapies.

HTN has been recognized as an inflammatory process, with even mild elevations in blood pressure associated with increased circulating IL-6 levels in otherwise healthy patients.^29,30^ Unfortunately, only about 30% of patients reach designated blood pressure goals through medical therapy, and evidence suggests that even those who successfully obtain normotension will still have vascular functional abnormalities.^31,32^ Notable consequences of inadequately treated HTN include aortic stiffening and left ventricular hypertrophy,^33^ both of which represent increased risk for cardiovascular morbidity and mortality.^34,35^ These tension-induced pathologies are driven by inflammatory cell infiltration with dysfunctional matrix remodeling,^36,37^ and indicate that interruption of cardiovascular mechanotransduction may represent a vital treatment pathway in HTN.

Blockade of SGK-1 activity has been shown to reduce fibrosis in other vascular beds by reduced signaling through NF-kB,^6,38,39^ a transcription factor known to influence expression of pro-inflammatory cytokines such as IL-6 and MCP-1,^40^ therefore our interest focused on SGK-1 as a mediator between mechanical aortic wall tension and inflammatory signaling. In this AngII-induced model of HTN in C57Bl/6 wild-type mice fed standard chow, aortic dissection and aneurysmal degeneration did not occur, but the elevated SBP did correspond to upregulated SGK-1 activity, increased circulating IL-6, and accumulation of macrophages within the abdominal aortic wall. Since macrophages are major producers of proteases,^41^ their abundance represents the potential for pathologic matrix remodeling. Interestingly, despite no change in blood pressure, treatment with EMD638683 reduced SGK-1 activity in parallel with decreased IL-6 levels and fewer macrophages, indicating that mechanical signaling was ongoing but the pathway toward inflammation was disrupted. The computational signaling network further supported this concept because inhibiting SGK-1 disconnected three major positive feedback loops that promote IL-6 expression. Targeting SGK-1 has significant therapeutic implications such that interrupting mechanical signaling as an adjunct to standard anti-hypertensive drugs may attenuate vascular remodeling to reduce end organ damage and warrants further investigation.

Because SGK-1 is ubiquitously expressed but the response to extracellular stress has been cell-type dependent,^6^ it was imperative to isolate murine abdominal aortic VSMCs to demonstrate by utilizing pharmacologic blockade, silencing RNA, and genetic knockout that this tension-induced cytokine production was SGK-1 dependent. Interestingly, despite the increase in MCP-1 expression and secretion in VSMCs under Stretch conditions, circulating levels of this cytokine were undetectable in the mice with AngII-induced HTN, raising a possibility that MCP-1 acted in a paracrine fashion within the aortic wall to promote macrophage accumulation. Alternatively, the production and secretion of IL-6 from aortic VSMCs likely represented a source for the elevated circulating IL-6 levels noted *in vivo*, and therefore mechanical activation of SGK-1 in aortic VSMCs promoted local and systemic inflammation. The computational network provided further evidence of integrated signaling by mechanical strain and AngII in aortic VSMCs with SGK-1 serving as a major hub for upregulation of IL-6 expression. As noted in the computational signaling network, NF-kB has been a known target for SGK-1 in VMSCs and can influence expression of both IL-6 and MCP-1.^42,43^ Interestingly, however, MCP-1 expression trended back to baseline with prolonged stimulation while IL-6 remained elevated, suggesting that negative feedback systems differentially impacted tension-induced cytokine expression. This observation corresponds to the pro- and anti-inflammatory effects reported for NF-kB and highlights the challenge of therapeutically targeting this transcription factor without undesirable side effects.^44^ With the concurrent evidence relating SGK-1 activity to VSMC proliferation, migration, and calcification based on alternative extracellular stressors such as glucose, adipokines, phosphates, and growth factors,^42,43,45–47^ integrating the impact of mechanical activation on inflammatory cell infiltration further demonstrates the value of pharmacologically inhibiting SGK-1 to abrogate vascular pathology.

### Limitations

Having demonstrated that tension-induced cytokine expression can be SGK-1 dependent and may represent a viable target to reduce dysfunctional hypertensive vascular remodeling, this project does have several limitations. First, while several prior studies have utilized salt-induced animal models of HTN to evaluate SGK-1, we chose to use the AngII-induced murine HTN model for the benefit of its parallelism with the human condition. This model will also allow subsequent integration of angiotensin converting enzyme inhibitors or angiotensin receptor blockers to determine how those medications influence SGK-1 activity and potential benefit for dual therapy with an SGK-1 inhibitor. Second, this project utilized a single *in vivo* HTN timepoint and cannot comment on the effect of longer term uncontrolled HTN or pharmacologically corrected SBP on aortic inflammation, but future studies may incorporate that experimental group. Third, having previously demonstrated that AngII exposure to Static murine aortic VSMCs did not significantly increase expression of IL-6 or MCP-1, and that concurrent treatment with AngII and Stretch had no additive effect over Stretch alone,^28^ we chose not to repeat all of those experiments with the SGK-1 inhibitor. This decision was further supported by the symmetry of the *in vitro* and *in vivo* model, but may be revisited in future studies utilizing anti-hypertensive medications to identify vital agents in VSMC mechanotransduction for SGK-1 activity. Finally, our computational modeling focused on the contributors to IL-6 expression since this was reflected in the *in vivo* murine experimentation, but future investigation may explore the signaling network specific to MCP-1 and its relationship to macrophage accumulation.

## Conclusion

In summary, mechanical activation of SGK-1 in aortic VSMCs can promote inflammatory signaling and macrophage infiltration, therefore this kinase warrants further exploration as a pharmacotherapeutic target to abrogate pathologic hypertensive vascular remodeling.

## Figure Legends

Supplement Figure 1. Representative flow cytometry gating to quantify CD11b^+^/F480^+^ cells.

Supplement Table 1. List of species integrated into the computational model.

Supplement Table 2. List of reactions integrated into the computational model.

