## Supplementary figures and images for "Vascular Smooth Muscle Cell Mechanotransduction Through Serum and Glucocorticoid Inducible Kinase -1 Promotes Interleukin-6 Production and Macrophage Accumulation in Murine Hypertension"

### Supplemental Fig 1

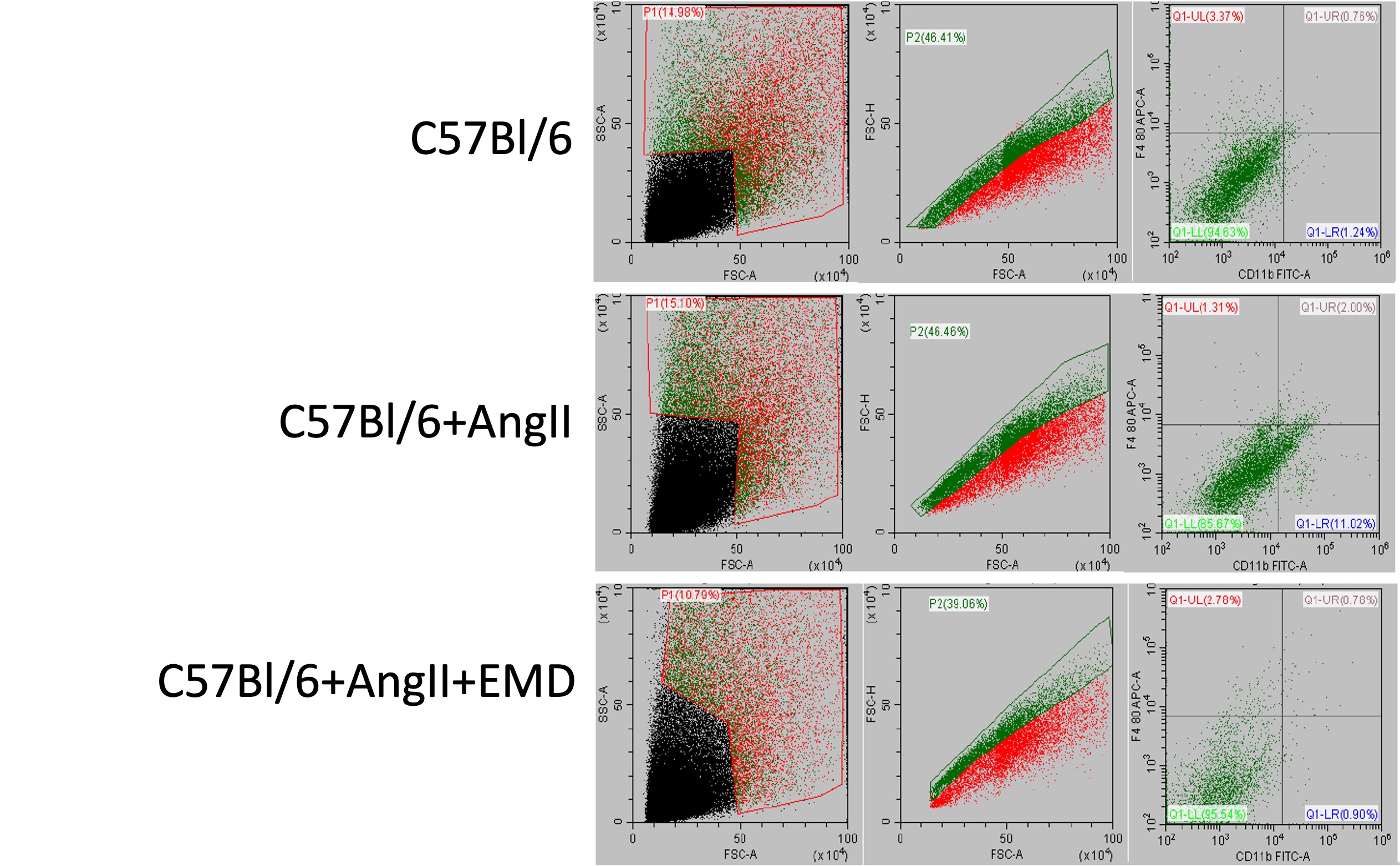
