## Supplementary material for "Vascular Smooth Muscle Cell Mechanotransduction Through Serum and Glucocorticoid Inducible Kinase -1 Promotes Interleukin-6 Production and Macrophage Accumulation in Murine Hypertension": Table 1

**Species information**

| module | ID | name | Yinit | Ymax | tau | type |
|---|---|---|---|---|---|---|
| macromolecule | Akt | Akt | 0 | 1 | 1 | protein |
| macromolecule | AMPK | AMPK | 0 | 1 | 1 | protein |
| macromolecule | AngII | AngiotensinII | 0 | 1 | 1 | protein |
| macromolecule | AngIIin | AngIIinput | 0 | 1 | 1 | protein |
| macromolecule | Apoptosis | Apoptosis | 0 | 1 | 1 | phenotype |
| macromolecule | AT1R | AngiotensinReceptor1 | 0 | 1 | 1 | protein |
| macromolecule | AT2R | AngiotensinReceptor2 | 0 | 1 | 1 | protein |
| macromolecule | betaCatenin | betaCatenin | 0 | 1 | 1 | protein |
| macromolecule | calponin | calponin | 0 | 1 | 1 | protein |
| macromolecule | Col3a1 | Col3a1 | 0 | 1 | 1 | mRNA |
| macromolecule | Collagen | Collagen | 0 | 1 | 1 | protein |
| macromolecule | DegElastin | DegradedElastin | 0 | 1 | 1 | protein |
| macromolecule | Dsh | Dsh | 0 | 1 | 1 | protein |
| macromolecule | Elastin | Elastin | 0 | 1 | 1 | protein |
| macromolecule | Elk1 | Elk1 | 0 | 1 | 1 | protein |
| macromolecule | Eln | Eln | 0 | 1 | 1 | mRNA |
| macromolecule | Energy | Energy | 0 | 1 | 1 | smallMolecule |
| macromolecule | ERK12 | ERK12 | 0 | 1 | 1 | protein |
| macromolecule | FAK | FAK | 0 | 1 | 1 | protein |
| macromolecule | Fibrillin | Fibrillin | 0 | 1 | 1 | protein |
| macromolecule | FOXO | FOXO | 0 | 1 | 1 | protein |
| macromolecule | Frizzled | Frizzled | 0 | 1 | 1 | protein |
| macromolecule | Glucose | Glucose | 0 | 1 | 1 | protein |
| macromolecule | GSK3 | GSK3 | 0 | 1 | 1 | protein |
| macromolecule | IGF | IGF | 0 | 1 | 1 | protein |
| macromolecule | IGFR | IGFR | 0 | 1 | 1 | protein |
| macromolecule | IL6 | IL6 | 0 | 1 | 1 | protein |
| macromolecule | IFNgamma | IFNgamma | 0 | 1 | 1 | protein |
| macromolecule | Integrins | Integrins | 0 | 1 | 1 | protein |
| macromolecule | IRS1 | IRS1 | 0 | 1 | 1 | protein |
| macromolecule | JAK | JAK | 0 | 1 | 1 | protein |
| macromolecule | JNK | JNK | 0 | 1 | 1 | protein |
| macromolecule | LAMP1 | LAMP1 | 0 | 1 | 1 | protein |
| macromolecule | LAMP2 | LAMP2 | 0 | 1 | 1 | protein |
| macromolecule | latTGFb | latentTGFbeta | 0 | 1 | 1 | protein |
| macromolecule | Leucine | Leucine | 0 | 1 | 1 | protein |
| macromolecule | MAPK | MAPK | 0 | 1 | 1 | protein |
| macromolecule | MEK | MEK | 0 | 1 | 1 | protein |
| macromolecule | mitf | *mitf* | 0 | 1 | 1 | mRNA |
| macromolecule | MITF | MITF | 0 | 1 | 1 | protein |
| macromolecule | MitoMet | MitoMetabolism | 0 | 1 | 1 | phenotype |
| macromolecule | MMP2 | MMP2 | 0 | 1 | 1 | protein |
| macromolecule | mTOR | mTOR | 0 | 1 | 1 | protein |
| macromolecule | mTORC1 | mTORC1 | 0 | 1 | 1 | protein |
| macromolecule | mTORC2 | mTORC2 | 0 | 1 | 1 | protein |
| macromolecule | NFkB | NFkappaB | 0 | 1 | 1 | protein |
| macromolecule | Oxygen | Oxygen | 0 | 1 | 1 | smallMolecule |
| macromolecule | p38 | p38 | 0 | 1 | 1 | protein |
| macromolecule | p4EBP1 | phosphorylated 4EBP1 | 0 | 1 | 1 | protein |
| macromolecule | PA | PA | 0 | 1 | 1 | protein |

| | | | | | | |
|---|---|---|---|---|---|---|
| macromolecule | PDGF | PDGFBB | 0 | 1 | 1 | protein |
| macromolecule | PDGFR | PDGFR | 0 | 1 | 1 | protein |
| macromolecule | PDK1 | PDK1 | 0 | 1 | 1 | protein |
| macromolecule | PI3K | PI3K | 0 | 1 | 1 | protein |
| macromolecule | PLD | PLD | 0 | 1 | 1 | protein |
| macromolecule | Raf | Raf | 0 | 1 | 1 | protein |
| macromolecule | Rag | Rag | 0 | 1 | 1 | protein |
| macromolecule | Ras | Ras | 0 | 1 | 1 | protein |
| macromolecule | RhoA | RhoA | 0 | 1 | 1 | protein |
| macromolecule | RSK1 | RSK1 | 0 | 1 | 1 | protein |
| macromolecule | S6 | S6 | 0 | 1 | 1 | protein |
| macromolecule | S6K | S6K1 | 0 | 1 | 1 | protein |
| macromolecule | Shear | Shear Stress | 0 | 1 | 1 | phenotype |
| macromolecule | SM22 | SM22 | 0 | 1 | 1 | protein |
| macromolecule | SMA | SMA | 0 | 1 | 1 | protein |
| macromolecule | Smad23 | Smad2/3 | 0 | 1 | 1 | protein |
| macromolecule | SMMHC | SMMHC | 0 | 1 | 1 | protein |
| macromolecule | SOCs | SOCs | 0 | 1 | 1 | protein |
| macromolecule | STAT | STAT | 0 | 1 | 1 | protein |
| phenotype | Stress | Intramural Stress | 0 | 1 | 1 | phenotype |
| phenotype | TGFbeta | TGFbeta | 0 | 1 | 1 | protein |
| phenotype | TGFR | TGFR | 0 | 1 | 1 | protein |
| phenotype | TIMP | TIMP | 0 | 1 | 1 | protein |
| macromolecule | TSC12 | TSC1/2 | 0 | 1 | 1 | protein |
| macromolecule | Wnt | WNT | 0 | 1 | 1 | protein |
| phenotype | Wnt5 | Wnt5 | 0 | 1 | 1 | protein |
| macromolecule | SGK1 | SGK1 | 0 | 1 | 1 | protein |
