## Supplementary material for "Vascular Smooth Muscle Cell Mechanotransduction Through Serum and Glucocorticoid Inducible Kinase -1 Promotes Interleukin-6 Production and Macrophage Accumulation in Murine Hypertension": Table 2

**Reaction Information**

| module | ID | Rule | Weight | n | EC50 | reflink 1 | reflink 2 | reflink 3 | reflink 4 | reflink 5 | reflink 6 | notes |
|---|---|---|---|---|---|---|---|---|---|---|---|---|
| inputs | i1 | => AngIIin | 0.1 | 1.4 | 0.52 | https://www.ahajournals.org/doi/abs/10.1161/01.hyp.29.1.366 | https://www.physiology.org/doi/10.1152/ajpheart.00040.2004 | | | | | |
| inputs | i2 | => Energy | 0.5 | 1.4 | 0.52 | https://science.sciencemag.org/content/294/5544/1102 | https://www.dovepress.com/recent-insights-into-the-pathophysiology-of-mtor-pathway-dysregulation-peer-reviewed-fulltext-article-RRB | | | | | |
| inputs | i3 | => Fibrillin | 0.25 | 1.4 | 0.52 | http://www.jbc.org/article/S0021925820837818/fulltext | | | | | | |
| inputs | i4 | => Glucose | 0.25 | 1.4 | 0.52 | https://www.dovepress.com/recent-insights-into-the-pathophysiology-of-mtor-pathway-dysregulation-peer-reviewed-fulltext-article-RRB | https://pubmed.ncbi.nlm.nih.gov/14651849/ | | | | | |
| inputs | i5 | => IFNgamma | 0.1 | 1.4 | 0.52 | https://www.nature.com/articles/375247a0 | | | | | | |
| inputs | i6 | => Leucine | 0.25 | 1.4 | 0.52 | https://www.dovepress.com/recent-insights-into-the-pathophysiology-of-mtor-pathway-dysregulation-peer-reviewed-fulltext-article-RRB | https://www.nature.com/articles/s41467-017-00785-0 | | | | | |
| inputs | i7 | => Oxygen | 0.5 | 1.4 | 0.52 | https://www.dovepress.com/recent-insights-into-the-pathophysiology-of-mtor-pathway-dysregulation-peer-reviewed-fulltext-article-RRB | http://genesdev.cshlp.org/content/18/23/2893.full | | | | | |
| inputs | i8 | => Stress | 0.1 | 1.4 | 0.52 | https://link.springer.com/article/10.1007/s12013-007-9002-3 | https://pubmed.ncbi.nlm.nih.gov/9588872/ | https://www.ahajournals.org/doi/abs/10.1161/atvbaha.107.147371 | https://www.ahajournals.org/doi/abs/10.1161/hh0701.089749 | https://journals.plos.org/plosone/article?id=10.1371/journal.pone.0070437 | https://faseb.onlinelibrary.wiley.com/doi/full/10.1096/fasebj.12.12.1135 | |
| middle | r1 | AngIIin => AngII | 1 | 1.4 | 0.52 | https://www.ahajournals.org/doi/abs/10.1161/01.hyp.29.1.366 | https://www.physiology.org/doi/10.1152/ajpheart.00040.2004 | | | | | |
| middle | r2 | AngII => AT1R | 1 | 1.4 | 0.52 | https://www.physiology.org/doi/10.1152/ajpheart.00040.2004 | | | | | | |
| middle | r3 | AngII => AT2R | 1 | 1.4 | 0.52 | https://www.physiology.org/doi/10.1152/ajpheart.00040.2004 | | | | | | |
| middle | r4 | Stress => AngII | 1 | 1.4 | 0.52 | https://pubmed.ncbi.nlm.nih.gov/9588872/ | | | | | | |
| middle | r5 | Stress => IGF | 1 | 1.4 | 0.52 | https://www.ahajournals.org/doi/abs/10.1161/atvbaha.107.147371 | | | | | | |
| middle | r6 | Stress => Integrins | 1 | 1.4 | 0.52 | https://www.ahajournals.org/doi/abs/10.1161/hh0701.089749 | | | | | | |
| middle | r7 | Stress => PDGF | 1 | 1.4 | 0.52 | https://journals.plos.org/plosone/article?id=10.1371/journal.pone.0070437 | https://faseb.onlinelibrary.wiley.com/doi/full/10.1096/fasebj.12.12.1135 | | | | | |
| middle | r8 | Stress => TGFbeta | 1 | 1.4 | 0.52 | https://pubmed.ncbi.nlm.nih.gov/9588872/ | | | | | | |
| middle | r9 | AT1R & JAK => PI3K | 1 | 1.4 | 0.52 | https://pubmed.ncbi.nlm.nih.gov/9242186/ | http://www.ncbi.nlm.nih.gov/pubmed/12623864 | | | | | |
| middle | r10 | AT1R => JAK | 1 | 1.4 | 0.52 | https://www.nature.com/articles/375247a0 | https://pubmed.ncbi.nlm.nih.gov/29876507/ | | | | | |
| middle | r11 | AT1R => JNK | 1 | 1.4 | 0.52 | https://www.ahajournals.org/doi/abs/10.1161/01.res.82.12.1272 | | | | | | |
| middle | r12 | AT1R => NFkB | 1 | 1.4 | 0.52 | https://www.ahajournals.org/doi/abs/10.1161/01.res.86.12.1266 | | | | | | |
| middle | r13 | AT1R => p38 | 1 | 1.4 | 0.52 | https://www.ncbi.nlm.nih.gov/pmc/articles/PMC3262055/ | | | | | | |
| middle | r14 | AT1R => PLD | 1 | 1.4 | 0.52 | https://pubmed.ncbi.nlm.nih.gov/29876507/ | https://pubmed.ncbi.nlm.nih.gov/8503884/ | https://mct.aacrjournals.org/content/11/5/1112 | | | | |
| middle | r15 | AT1R => Ras | 1 | 1.4 | 0.52 | http://www.jbc.org/article/S0021925818467816/fulltext | | | | | | |
| middle | r16 | AT1R => Smad23 | 1 | 1.4 | 0.52 | http://www.ncbi.nlm.nih.gov/pubmed/23868934 | | | | | | |
| middle | r17 | AT1R => TIMP | 1 | 1.4 | 0.52 | https://pubmed.ncbi.nlm.nih.gov/12388255/ | | | | | | |
| middle | r18 | AT1R => Wnt5 | 1 | 1.4 | 0.52 | https://www.nature.com/articles/s41598-018-27064-2 | | | | | | |
| middle | r19 | !Energy & !Oxygen & AT1R => AMPK | 1 | 1.4 | 0.52 | https://pubmed.ncbi.nlm.nih.gov/16959574/ | | | | | | |
| middle | r20 | latTGFb & !Fibrillin => TGFbeta | 1 | 1.4 | 0.52 | https://onlinelibrary.wiley.com/doi/full/10.1002/jcp.24094 | | | | | | |
| middle | r21 | MEK & !AT2R => ERK12 | 1 | 1.4 | 0.52 | https://www.ahajournals.org/doi/abs/10.1161/atvbaha.109.200501 | https://science.sciencemag.org/content/332/6027/361 | | | | | |
| middle | r22 | mTORC2 & !AT2R => RhoA | 1 | 1.4 | 0.52 | https://pubmed.ncbi.nlm.nih.gov/15467718/ | https://pubmed.ncbi.nlm.nih.gov/17082724/ | | | | | |
| middle | r23 | !Akt & !Dsh => GSK3 | 1 | 1.4 | 0.52 | https://pubmed.ncbi.nlm.nih.gov/11437438/ | https://pubmed.ncbi.nlm.nih.gov/17052453/ | | | | | |
| middle | r24 | !Akt & !SGK1 => FOXO | 1 | 1.4 | 0.52 | https://pubmed.ncbi.nlm.nih.gov/17052453/ | https://pubmed.ncbi.nlm.nih.gov/15380067/ | doi:10.1177/1758835920940946 | | | | |
| middle | r25 | !S6K => IRS1 | 1 | 1.4 | 0.52 | https://pubmed.ncbi.nlm.nih.gov/15380067/ | | | | | | |
| middle | r26 | !Dsh & !S6K => GSK3 | 1 | 1.4 | 0.52 | https://pubmed.ncbi.nlm.nih.gov/11437438/ | https://pubmed.ncbi.nlm.nih.gov/17052453/ | | | | | |
| middle | r27 | !GSK3 => betaCatenin | 1 | 1.4 | 0.52 | https://pubmed.ncbi.nlm.nih.gov/11437438/ | | | | | | |
| middle | r28 | !TSC12 => mTORC1 | 1 | 1.4 | 0.52 | https://pubmed.ncbi.nlm.nih.gov/15380067/ | https://pubmed.ncbi.nlm.nih.gov/16959574/ | | | | | |
| middle | r29 | AMPK & !Akt & !ERK12 & !p38 & !RSK1 => TSC12 | 1 | 1.4 | 0.52 | https://pubmed.ncbi.nlm.nih.gov/16959574/ | https://www.nature.com/articles/ncb839 | https://pubmed.ncbi.nlm.nih.gov/15851026/ | https://pubmed.ncbi.nlm.nih.gov/12582162/ | https://pubmed.ncbi.nlm.nih.gov/15342917/ | https://pubmed.ncbi.nlm.nih.gov/12150915/ | https://pubmed.ncbi.nlm.nih.gov/17908691/ |
| middle | r30 | GSK3 & !Akt & !ERK12 & !p38 & !RSK1 => TSC12 | 1 | 1.4 | 0.52 | https://pubmed.ncbi.nlm.nih.gov/16959574/ | https://www.nature.com/articles/ncb839 | https://pubmed.ncbi.nlm.nih.gov/15851026/ | https://pubmed.ncbi.nlm.nih.gov/12582162/ | https://pubmed.ncbi.nlm.nih.gov/15342917/ | https://pubmed.ncbi.nlm.nih.gov/12150915/ | https://pubmed.ncbi.nlm.nih.gov/17908691/ |
| middle | r31 | SGK1 & !Akt & !ERK12 & !p38 & !RSK1 => TSC12 | 1 | 1.4 | 0.52 | doi:10.1177/1758835920940946 | https://www.nature.com/articles/ncb839 | https://pubmed.ncbi.nlm.nih.gov/15851026/ | https://pubmed.ncbi.nlm.nih.gov/12582162/ | https://pubmed.ncbi.nlm.nih.gov/15342917/ | https://pubmed.ncbi.nlm.nih.gov/12150915/ | https://pubmed.ncbi.nlm.nih.gov/17908691/ |
| middle | r32 | AMPK => JNK | 1 | 1.4 | 0.52 | https://pubmed.ncbi.nlm.nih.gov/19037093/ | | | | | | |
| middle | r33 | ERK12 => Elk1 | 1 | 1.4 | 0.52 | https://www.ahajournals.org/doi/abs/10.1161/01.HYP.0000054213.37471.84 | | | | | | |
| middle | r34 | ERK12 => RSK1 | 1 | 1.4 | 0.52 | https://journals.physiology.org/doi/abs/10.1152/ajpcell.00552.2005 | | | | | | |
| middle | r35 | FAK => JNK | 1 | 1.4 | 0.52 | https://pubmed.ncbi.nlm.nih.gov/12782622/ | | | | | | |
| middle | r36 | FAK => Ras | 1 | 1.4 | 0.52 | https://pubmed.ncbi.nlm.nih.gov/7997267/ | | | | | | |
| middle | r37 | Fibrillin => Integrins | 1 | 1.4 | 0.52 | http://www.jbc.org/article/S0021925820837818/fulltext | | | | | | |
| middle | r38 | Frizzled => Dsh | 1 | 1.4 | 0.52 | https://pubmed.ncbi.nlm.nih.gov/11437438/ | | | | | | |
| middle | r39 | Glucose => MitoMet | 1 | 1.4 | 0.52 | https://science.sciencemag.org/content/294/5 | | | | | | |

| | | | | | | | | |
|---|---|---|---|---|---|---|---|---|
| middle | r40 | IGF => Eln | 1 | 1.4 | 0.52 | https://pubmed.ncbi.nlm.nih.gov/8509381/ | | |
| middle | r41 | IGF => IGFR | 1 | 1.4 | 0.52 | https://pubmed.ncbi.nlm.nih.gov/17908691/ | | |
| middle | r42 | IGFR => Ras | 1 | 1.4 | 0.52 | https://pubmed.ncbi.nlm.nih.gov/12943991/ | | |
| middle | r43 | IL6 & !SOCs => JAK | 1 | 1.4 | 0.52 | https://europepmc.org/article/med/15256805 | https://pubmed.ncbi.nlm.nih.gov/12150892/ | |
| middle | r44 | IFNgamma => JAK | 1 | 1.4 | 0.52 | https://www.nature.com/articles/375247a0 | | |
| middle | r45 | Integrins => FAK | 1 | 1.4 | 0.52 | https://www.ahajournals.org/doi/abs/10.1161/01.cir.0000154548.16191.2f | | |
| middle | r46 | Integrins => latTGFb | 1 | 1.4 | 0.52 | https://pubmed.ncbi.nlm.nih.gov/25670798/ | | |
| middle | r47 | IRS1 & IGFR => PI3K | 1 | 1.4 | 0.52 | https://pubmed.ncbi.nlm.nih.gov/15380067/ | https://pubmed.ncbi.nlm.nih.gov/17908691/ | |
| middle | r48 | JAK => STAT | 1 | 1.4 | 0.52 | https://www.nature.com/articles/375247a0 | | |
| middle | r49 | Leucine => Rag | 1 | 1.4 | 0.52 | https://www.nature.com/articles/s41467-017-00785-0 | | |
| middle | r50 | MAPK => latTGFb | 1 | 1.4 | 0.52 | https://journals.physiology.org/doi/abs/10.1152/ajpcell.00008.2012 | https://pubmed.ncbi.nlm.nih.gov/16624629/ | |
| middle | r51 | MAPK => MEK | 1 | 1.4 | 0.52 | https://www.ahajournals.org/doi/abs/10.1161/atvbaha.109.200501 | | |
| middle | r52 | mitf & GSK3 & mTORC1 => MITF | 1 | 1.4 | 0.52 | https://pubmed.ncbi.nlm.nih.gov/30150413/ | | |
| middle | r53 | Akt => mTOR | 1 | 1.4 | 0.52 | https://www.ahajournals.org/doi/abs/10.1161/atvbaha.108.179457 | | |
| middle | r54 | MitoMet & mTOR => mTORC1 | 1 | 1.4 | 0.52 | https://science.sciencemag.org/content/294/5544/1102 | | |
| middle | r55 | mTOR => mTORC2 | 1 | 1.4 | 0.52 | https://www.nature.com/articles/s41598-019-56237-w | | |
| middle | r56 | mTORC1 => p4EBP1 | 1 | 1.4 | 0.52 | https://pubmed.ncbi.nlm.nih.gov/14592809/ | | |
| middle | r57 | mTORC1 => S6K | 1 | 1.4 | 0.52 | https://pubmed.ncbi.nlm.nih.gov/14592809/ | | |
| middle | r58 | mTORC2 => Akt | 1 | 1.4 | 0.52 | https://www.nature.com/articles/s41598-019-56237-w | https://faseb.onlinelibrary.wiley.com/doi/full/10.1096/fj.10-175018 | |
| middle | r59 | NFkB => IL6 | 1 | 1.4 | 0.52 | https://journals.physiology.org/doi/abs/10.1152/ajpheart.00919.2004 | | |
| middle | r60 | NFkB => Wnt5 | 1 | 1.4 | 0.52 | https://pubmed.ncbi.nlm.nih.gov/19424602/ | | |
| middle | r61 | p38 => Col3a1 | 1 | 1.4 | 0.52 | https://pubmed.ncbi.nlm.nih.gov/11230337/ | | |
| middle | r62 | PA & mTOR => mTORC1 | 1 | 1.4 | 0.52 | https://pubmed.ncbi.nlm.nih.gov/16537399/ | https://pubmed.ncbi.nlm.nih.gov/23077579/ | |
| middle | r63 | PDGF => PDGFR | 1 | 1.4 | 0.52 | https://journals.plos.org/plosone/article?id=10.1371/journal.pone.0070437 | | |
| middle | r64 | PDGFR => JNK | 1 | 1.4 | 0.52 | https://faseb.onlinelibrary.wiley.com/doi/full/10.1096/fasebj.12.12.1135 | https://pubmed.ncbi.nlm.nih.gov/16854986/ | |
| middle | r65 | PDGFR => p38 | 1 | 1.4 | 0.52 | https://www.ahajournals.org/doi/abs/10.1161/ATVBAHA.116.308895 | | |
| middle | r66 | PDGFR => Ras | 1 | 1.4 | 0.52 | https://pubmed.ncbi.nlm.nih.gov/2156626/ | https://journals.plos.org/plosbiology/article?id=10.1371/journal.pbio.0000052 | |
| middle | r67 | PDGFR=> PI3K | 1 | 1.4 | 0.52 | https://journals.plos.org/plosone/article?id=10.1371/journal.pone.0070437 | | |
| middle | r68 | PDK1 => Akt | 1 | 1.4 | 0.52 | http://www.ncbi.nlm.nih.gov/pubmed/12623864 | | |
| middle | r69 | PI3K => PDK1 | 1 | 1.4 | 0.52 | http://www.ncbi.nlm.nih.gov/pubmed/12623864 | | |
| middle | r70 | PLD => PA | 1 | 1.4 | 0.52 | https://pubmed.ncbi.nlm.nih.gov/16537399/ | | |
| middle | r71 | Raf => MAPK | 1 | 1.4 | 0.52 | https://www.ahajournals.org/doi/abs/10.1161/atvbaha.109.200501 | | |
| middle | r72 | Rag & mTOR => mTORC1 | 1 | 1.4 | 0.52 | https://www.nature.com/articles/s41467-017-00785-0 | https://www.nature.com/articles/ncb1753 | |
| middle | r73 | Ras => Raf | 1 | 1.4 | 0.52 | https://www.ahajournals.org/doi/abs/10.1161/atvbaha.109.200501 | | |
| middle | r74 | Ras => RhoA | 1 | 1.4 | 0.52 | https://pubmed.ncbi.nlm.nih.gov/10400905/ | | |
| middle | r75 | S6K => S6 | 1 | 1.4 | 0.52 | https://pubmed.ncbi.nlm.nih.gov/17052453/ | https://pubmed.ncbi.nlm.nih.gov/15380067/ | |
| middle | r76 | Smad23 => Col3a1 | 1 | 1.4 | 0.52 | https://www.nature.com/articles/srep19503 | https://pubmed.ncbi.nlm.nih.gov/23091366/ | |
| middle | r77 | Smad23 => Eln | 1 | 1.4 | 0.52 | https://pubmed.ncbi.nlm.nih.gov/32404006/ | | |
| middle | r78 | STAT => SOCs | 1 | 1.4 | 0.52 | https://pubmed.ncbi.nlm.nih.gov/12150892/ | https://pubmed.ncbi.nlm.nih.gov/18708154/ | |
| middle | r79 | STAT => Wnt5 | 1 | 1.4 | 0.52 | https://pubmed.ncbi.nlm.nih.gov/19424602/ | | |
| middle | r80 | TGFbeta => TGFR | 1 | 1.4 | 0.52 | https://pubmed.ncbi.nlm.nih.gov/9588872/ | | |
| middle | r81 | TGFR & TSC12 => Smad23 | 1 | 1.4 | 0.52 | http://www.jbc.org/article/S0021925820883641/fulltext | https://pubmed.ncbi.nlm.nih.gov/25727005/ | https://jpet.aspetjournals.org/content/315/3/1005 |
| middle | r82 | TGFR => p38 | 1 | 1.4 | 0.52 | https://jpet.aspetjournals.org/content/315/3/1005 | | |
| middle | r83 | TGFR => TIMP | 1 | 1.4 | 0.52 | https://journals.plos.org/plosone/article?id=10.1371/journal.pone.0014145 | | |
| middle | r84 | TGFR => Wnt5 | 1 | 1.4 | 0.52 | https://pubmed.ncbi.nlm.nih.gov/19424602/ | | |
| middle | r85 | TGFR=> JNK | 1 | 1.4 | 0.52 | https://jpet.aspetjournals.org/content/315/3/1005 | | |
| middle | r86 | Wnt => Frizzled | 1 | 1.4 | 0.52 | https://pubmed.ncbi.nlm.nih.gov/11437438/ | | |
| middle | r87 | Wnt5 => Wnt | 1 | 1.4 | 0.52 | https://pubmed.ncbi.nlm.nih.gov/19424602/ | | |
| middle | r88 | PDK1 => SGK1 | 1 | 1.4 | 0.52 | doi:10.1177/1758835920940946 | | |
| middle | r89 | mTORC2 => SGK1 | 1 | 1.4 | 0.52 | doi:10.1177/1758835920940946 | | |
| middle | r90 | SGK1 => NFkB | 1 | 1.4 | 0.52 | doi:10.1177/1758835920940946 | | |
| outputs | r91 | !Elk1 & RhoA => calponin | 1 | 1.4 | 0.52 | https://www.nature.com/articles/nature02382 | | |

| | | | | | | | | | | |
|---|---|---|---|---|---|---|---|---|---|---|
| outputs | r92 | !Elk1 & RhoA => SM22 | 1 | 1.4 | 0.52 | https://www.nature.com/articles/nature02382 | | | | |
| outputs | r93 | !Elk1 & RhoA => SMA | 1 | 1.4 | 0.52 | https://www.nature.com/articles/nature02382 | | | | |
| outputs | r94 | !Elk1 & RhoA => SMMHC | 1 | 1.4 | 0.52 | https://www.nature.com/articles/nature02382 | | | | |
| outputs | r95 | betaCatenin => MMP2 | 1 | 1.4 | 0.52 | https://www.ahajournals.org/doi/10.1161/ATVBAHA.116.308643 | | | | |
| outputs | r96 | betaCatenin => LAMP1 | 1 | 1.4 | 0.52 | https://www.pnas.org/content/116/21/10402 | https://pubmed.ncbi.nlm.nih.gov/32726636/ | | | |
| outputs | r97 | betaCatenin => LAMP2 | 1 | 1.4 | 0.52 | https://www.pnas.org/content/116/21/10402 | https://pubmed.ncbi.nlm.nih.gov/32726636/ | | | |
| outputs | r98 | betaCatenin => mitf | 1 | 1.4 | 0.52 | https://pubmed.ncbi.nlm.nih.gov/27918305/ | | | | |
| outputs | r99 | Col3a1 => Collagen | 1 | 1.4 | 0.52 | https://www.pnas.org/content/94/5/1852 | | | | |
| outputs | r100 | Eln & Fibrillin => Elastin | 1 | 1.4 | 0.52 | https://pubmed.ncbi.nlm.nih.gov/31395654/ | | | | |
| outputs | r101 | ERK12 => MMP2 | 1 | 1.4 | 0.52 | https://pubmed.ncbi.nlm.nih.gov/16854986/ | https://pubmed.ncbi.nlm.nih.gov/24792035/ | | | |
| outputs | r102 | ERK12 => SMMHC | 1 | 1.4 | 0.52 | https://journals.plos.org/plosone/article?id=10.1371/journal.pone.0012196 | https://www.ahajournals.org/doi/abs/10.1161/01.HYP.0000054213.37471.84 | | | |
| outputs | r103 | FOXO => Apoptosis | 1 | 1.4 | 0.52 | https://pubmed.ncbi.nlm.nih.gov/15380067/ | | | | |
| outputs | r104 | MITF => LAMP1 | 1 | 1.4 | 0.52 | https://www.pnas.org/content/112/5/E420 | https://www.jci.org/articles/view/128287 | | | |
| outputs | r105 | MITF => LAMP2 | 1 | 1.4 | 0.52 | https://www.jci.org/articles/view/128287 | https://journals.plos.org/plosone/article?id=10.1371/journal.pone.0173771 | | | |
| outputs | r106 | Smad23 & p38 & !FOXO & !NFkB => SMA | 1 | 1.4 | 0.52 | http://www.cell.com/article/S1534580705002133/fulltext | https://www.pnas.org/content/105/9/3362 | https://pubmed.ncbi.nlm.nih.gov/21832838/ | https://www.ncbi.nlm.nih.gov/pmc/articles/PMC3442374/ | |
| outputs | r107 | Smad23 & p38 & !FOXO & !NFkB => calponin | 1 | 1.4 | 0.52 | http://www.cell.com/article/S1534580705002133/fulltext | https://www.pnas.org/content/105/9/3362 | https://pubmed.ncbi.nlm.nih.gov/21832838/ | https://www.ncbi.nlm.nih.gov/pmc/articles/PMC3442374/ | |
| outputs | r108 | Smad23 & p38 & !FOXO & !NFkB => SM22 | 1 | 1.4 | 0.52 | http://www.cell.com/article/S1534580705002133/fulltext | https://www.pnas.org/content/105/9/3362 | https://pubmed.ncbi.nlm.nih.gov/21832838/ | https://www.ncbi.nlm.nih.gov/pmc/articles/PMC3442374/ | |
| outputs | r109 | Smad23 & p38 & !FOXO & !NFkB => SMMHC | 1 | 1.4 | 0.52 | http://www.cell.com/article/S1534580705002133/fulltext | https://www.pnas.org/content/105/9/3362 | https://www.ncbi.nlm.nih.gov/pmc/articles/PMC3442374/ | | |
| outputs | r110 | Akt & !FOXO & !NFkB => calponin | 1 | 1.4 | 0.52 | https://pubmed.ncbi.nlm.nih.gov/17908691/ | https://www.ahajournals.org/doi/abs/10.1161/atvbaha.108.179457 | https://pubmed.ncbi.nlm.nih.gov/25201081/ | http://www.cell.com/article/S1534580705002133/fulltext | https://www.pnas.org/content/105/9/3362 |
| outputs | r111 | Akt & !FOXO & !NFkB => SM22 | 1 | 1.4 | 0.52 | https://pubmed.ncbi.nlm.nih.gov/17908691/ | https://www.ahajournals.org/doi/abs/10.1161/atvbaha.108.179457 | https://pubmed.ncbi.nlm.nih.gov/25201081/ | http://www.cell.com/article/S1534580705002133/fulltext | https://www.pnas.org/content/105/9/3362 |
| outputs | r112 | Akt & !FOXO & !NFkB => SMA | 1 | 1.4 | 0.52 | https://pubmed.ncbi.nlm.nih.gov/17908691/ | https://www.ahajournals.org/doi/abs/10.1161/atvbaha.108.179457 | https://pubmed.ncbi.nlm.nih.gov/25201081/ | http://www.cell.com/article/S1534580705002133/fulltext | https://www.pnas.org/content/105/9/3362 |
| outputs | r113 | Akt & !FOXO & !NFkB => SMMHC | 1 | 1.4 | 0.52 | https://pubmed.ncbi.nlm.nih.gov/17908691/ | https://www.ahajournals.org/doi/abs/10.1161/atvbaha.108.179457 | http://www.cell.com/article/S1534580705002133/fulltext | https://www.pnas.org/content/105/9/3362 | |
| outputs | r114 | Akt & JNK & p38 & !TIMP & !TSC12 => MMP2 | 1 | 1.4 | 0.52 | https://journals.plos.org/plosone/article?id=10.1371/journal.pone.0070437 | https://pubmed.ncbi.nlm.nih.gov/16854986/ | https://pubmed.ncbi.nlm.nih.gov/24792035/ | | |
| outputs | r115 | Akt => TIMP | 1 | 1.4 | 0.52 | https://www.ahajournals.org/doi/abs/10.1161/ATVBAHA.117.310502 | | | | |
